# Selecting Reads for Haplotype Assembly

**DOI:** 10.1101/046771

**Authors:** Sarah O. Fischer, Tobias Marschall

**Affiliations:** Center for Bioinformatics, Saarland University, Saarbrücken, Germany; Max Planck Institute for Informatics, Saarbrücken, Germany

## Abstract

*Haplotype assembly or read-based phasing* is the problem of reconstructing both haplotypes of a diploid genome from next-generation sequencing data. This problem is formalized as the Minimum Error Correction (MEC) problem and can be solved using algorithms such as WhatsHap. The runtime of WhatsHap is exponential in the maximum coverage, which is hence controlled in a pre-processing step that selects reads to be used for phasing. Here, we report on a heuristic algorithm designed to choose beneficial reads for phasing, in particular to increase the connectivity of the phased blocks and the number of correctly phased variants compared to the random selection previously employed in by WhatsHap. The algorithm we describe has been integrated into the WhatsHap software, which is available under MIT licence from https://bitbucket.org/whatshap/whatshap.

## 1 Introduction

In diploid species, mother and father each pass on one copy of every chromosome to their offspring. The task of reconstructing these two chromosomal sequences, which are called haplotypes, is known as *phasing* or *haplotyping* (Tewhey *et al.*, 2011; Glusman *et al.*, 2014). Next-generation sequencing (NGS) reads that are sufficiently long to cover two or more heterozygous variants are *phase informative* and can be used for this purpose. The computational problem of inferring the two haplotypes from (aligned) NGS data is known as *read-based phasing* or *haplotype assembly*. Its most common and most successful formalization is the Minimum Error Correction (MEC) problem, which is NP-hard (Cilibrasi *et al.*, 2007). Among others, the ideas of fixed-parameter tractability (FPT) have been applied to attack this problem (He *et al.*, 2010; Patterson *et al.*, 2015).

The runtime of the WhatsHap algorithm (Patterson *et al.*, 2015) is exponential in the maximum coverage but only linear in the number of phased variants and independent of the read length. These properties make it particularly suited for long-read data (such as delivered by PacBio or Oxford Nanopore devices). However, the exponential runtime in the maximum coverage requires the preprocessing step of ensuring this quantity to be bounded. This is achieved by discarding reads in regions of excess coverage. Patterson *et al.* (2015) use a user-specified parameter for the maximum coverage and select reads in a random way: the reads are enumerated in random order and each read is retained if it is phase informative and adding it does not violate the coverage constraint (given the previously selected reads). Figure 1 illustrates that such a random selection can lead to undesirable results.

**Figure 1:**
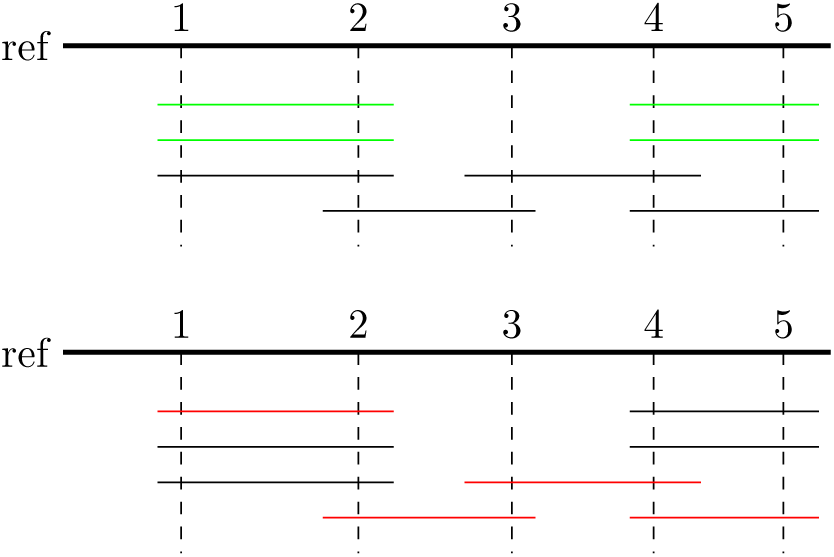
Example of selecting reads (horizontal lines) that cover five different SNPs (indicated by dashed vertical lines) with a maximum coverage of two. **Top:** unfortunate selection (green); although no more reads can be added without violating the coverage constraint, SNP 3 is not covered at all and SNPs 1 and 2 are not connected to SNPs 3 and 4. **Bottom:** better selection (red) that covers and connects all SNPs.

In this paper, we propose an alternative algorithm to select reads under a given coverage constraint. It is a greedy heuristic that aims to exhibit the following desirable properties:

1. as many heterozygous variants as possible should be covered,
2. the covered variants should be covered by as many reads as possible,
3. reads covering many variants at once should be preferred,
4. high-quality reads (in terms of mapping and base-calling quality) shall be preferred over low quality ones,
5. all variants should be well connected by reads, i.e. the number of connected components in the resulting graph (nodes: variants, edges: two variants covered by a selected read) should be low, and
6. each pair of variants should be independently connected by different paths as often as possible.

Many different formalizations for the *read selection problem* based on these desirable properties are conceivable. How to best trade-off these partly conflicting properties is an open research question and little literature on it exists. Mäkinen *et al.* (2015) propose to maximize the minimum coverage by means of a flow-based approach. In the following, we introduce a heuristic algorithm that we show to work well in practice. That is, we demonstrate that haplotype assembly performed on the selected reads yields good results.

## 2 Read Selection Algorithm

As a first step, all reads covering less than two heterozygous variants can be discarded since they are not phase informative. In the following, we thus assume all reads to cover at least two heterozygous variants.

Our algorithms works iteratively. In each iteration, a subset of reads is selected, which we call a *slice*. All slices are disjoint, that is, reads already part of a slice are not considered in later iterations. Each slice tries to cover all variants (Goal 1 in Section 1) and to connect as many variants as possible (Goal 5), while using as few reads as possible. Figure 2 illustrates that each slice could archieve these goals. Therefore, every individual slice connects (in the best case) all variants to each other and hence provides a connection between each pair of variants which is independent of the other slices, catering to Goal 6.

**Figure 2:**
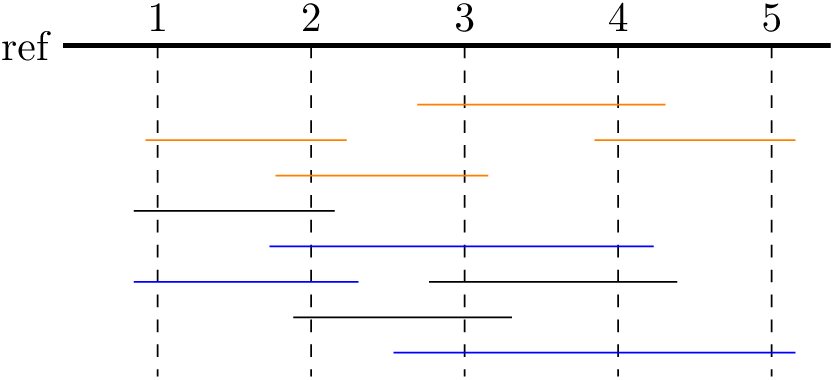
Example of two slices (indicated by coloured reads) that cover five different SNPs with a maximum coverage of four. Every slice covers every SNP and set up different connectivity patterns between the five SNPs.

Each iteration, i.e. selecting reads for a slice, consists of two phases, which both use a *score* measuring the “usefulness” of a read (detailed in Section 2.1 below): First, reads are enumerated ordered by score and those that cover at least one variant thus far uncovered (in the present slice) are greedily added. Second, reads *bridging* two connected components within that slice are added, again greedily in the order induced by the score. Before adding a read (in either of the two steps), we test whether doing so would violate the coverage constraints and, if so, discard it.

These two steps are repeated to add slice after slice until no further reads are left.

### 2.1 Scoring

We introduce a scoring function that intends to select reads that cover as many variants as possible (Goal 3) and have a high quality (Goal 4). Paired-end or mate-pair reads can cover variants that are not consecutive. We call uncovered variants that lie between covered variants (for a given read pair) *holes*. Selecting read pairs with holes is undesirable because holes contribute to the (physical) coverage at a particular variant, but do not any information.

To define the scoring function, we introduce the following notation. Let 𝓡 be the set of all reads. For *R* ∈ 𝓡, let `variants`(*R*) and holes(*R*) denote the set of variants covered by *R* and the set of holes of *R*, respectively. Furthermore, quality(*R*, *V*) denotes the base quality of the nucleotide in read *R* covering variant *V* ∈ `variants`(*R*). By 𝓡_*s*_ ⊂ 𝓡, we refer to the set of reads already selected for the current slice.

We define three different scores for a read *R*. The first one is defined through

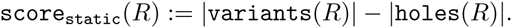

It is called `score`_static_ because its value does not change in the course of the algorithm. In contrast, `score`_dyn_ changes as reads get added to a slice:

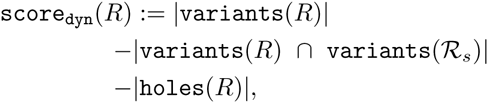

where variants(𝓡_*s*_) refers to the set of all variants covered by reads in 𝓡_*s*_, formally

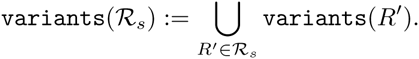

Therefore, `score`_dyn_(*R*) is the number of variants covered by *R* that are not yet covered by any read in 𝓡_*s*_ minus the number of holes. It is thus useful to assess the value of adding *R* to slice 𝓡_*s*_. The third score we consider is defined as

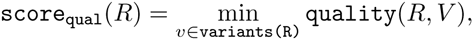

that is, it gives the quality value of the variant covered by that read with worst quality. To rank reads, we compare them by the tuple score

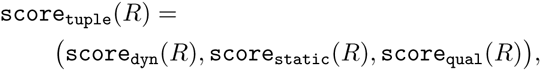

that is, two reads are first compared by means of `score`_dyn_, then (in case of a tie) by `score`_static_, and as a last ressort by `score`_qual_.

#### Algorithm 1 Score-based read selection

1: **procedure** ReadSelection(`readset, max_cov`)

2: `selected_reads` ← empty set

3: `undecided_reads` ← empty set

4: **for** *R* in `readset` **do**

5: **if** |`variants`(*R*)| ≥ 2 **then**

6: **undecided_reads**.Add(*R*)

7: **end if**

8: **end for**

9: **while** `undecided_reads` not empty **do**

10: (`reads_in_slice, reads_violating_coverage`) ← SelectSlice(`undecided_reads, max_cov`)

11: `selected_reads`.Add(`reads_in_slice`)

12: `undecided_reads`.Remove(`reads_in_slice`)

13: `undecided_reads`.Remove(`reads_violating_coverage`)

14: `bridge_reads` ← Bridging(`reads_in_slice, undecided_reads, max_cov`) ▹ Optional bridging step

15: `selected_reads`.Add(`bridge_reads`) ▹ Optional bridging step

16: **end while**

17: **return** `selected_reads`

18: **end procedure**

### 2.2 Algorithm

Pseudo code of our read selection algorithm is given as Algorithm 1. At first, all reads that cover at least two heterozygous variants are stored in the set `undecided_reads` (lines 4 to 8). In the course of the algorithm they are moved to `selected_reads` or discarded. Each iteration of the while loop in Line 9 creates one slice by calling SelectSlice and Bridging and terminates when no undecided reads are left.

In SelectSlice (see Algorithm 2), a priority queue is constructed from `undecided_reads`, using `score`_tuple_ as a scoring function. Based on this priority queue a set of reads is selected, extracting the best reads one after each other until every variant is covered once or no usable reads are left. This function maintains a set `already_covered_snps` with variants covered by any read selected so far. Based on this set, the variants additionally covered by this read are determined (`snps_covered_by_this_read`). Only reads for which this set is non-empty and which do not violate the coverage constraint are selected and added to `reads_in_slice`. Since `score`_dyn_ of a read depends on the reads that have already been selected in a slice, we need to update these scores. Adding a read can lead to changed scores for other reads that cover the same SNPs, while not affecting reads that cover a disjoint set of variants. In lines 19 to 27 of Algorithm 2, the set of reads to be updated is determined, the scores recomputed and updated accordingly in the priority queue. Note that this requires an extra index that maps variants to all reads covering them (which is not explicitly mentioned in the pseudo code).

The function Bridging given in Algorithm 3 is called by Algorithm 1 (in Line 14) to add reads that can lower the number of connected components and hence increase connectivity. Again, reads are enumerated ordered by score. A union-find data structure Cormen *et al.* (2009) is used to determine whether a read connects two components and, in case it does, the read is greedily added. Figure 3 illustrates this bridging step.

**Figure 3:**
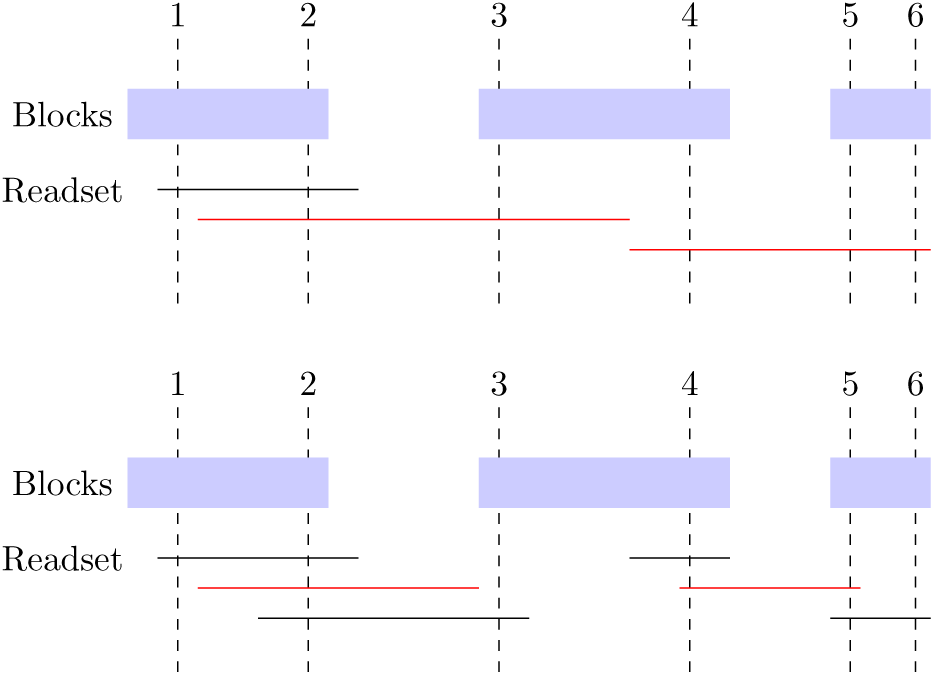
Illustration of reads, selected in the bridging step of the scoring based read selection. The blue blocks indicate connected components of reads selected previously (in Algorithm 2); the horizontal lines represent yet undecided reads. Reads highlighted in red are selected because they connect previously unconnected blocks. **Top:** single-end reads. **Bottom:** two lines in one row represent a paired-end read, i.e. there is phase information between the two reads in a pair.

## 3 Evaluation

The evaluation of our score-based read selection is based on the comparison of this approach with the random approach. We generated (simulated) benchmark data sets using the same procedure as for evaluation presented by Patterson *et al.* (2015). Furthermore, we added a variant of our approach that omits the bridging step. We ran the three read selection methods to generated data sets with 5× and 15× target coverage. After read selection, the pruned read sets are phased using

### Algorithm 2 Select one slice of reads

1: **procedure** SelectSlice(`undecided_reads, max_cov`)

2: `already_covered_snps` ← empty set

3: `reads_in_slice` ← empty set

4: `reads_violating_coverage` ← empty set

5: *pq* ← ConstructPriorityqueue(`undecided_reads`)

6: **while** *pq* is not empty **do**

7: `snps_covered_by_this_read` ← empty set

8: (`score, read`) ← *pq*.Pop

9: **for** *V* in variants(`read`) **do**

10: **if** V.*pos* not in `already_covered_snps` **then**

11: `snps_covered_by_this_read`.Add(*V.pos*)

12: **end if**

13: **end for**

14: **if** adding read would `exceed max_cov` for at least one position **then**

15: `reads_violating_coverage`.Add(read)

16: **else**

17: **if** `snps_covered_by_this_read` not empty **then**

18: `reads_in_slice`.Add(read)

19: `reads_to_be_updated` ← empty set

20: **for** *pos* in `snps_covered_by_this_read` **do**

21: `already_covered_snps`.Add(*pos*)

22: `reads_to_be_updated`.Add(all reads in *pq* covering *pos*)

23: **end for**

24: **for** *R* in `reads_to_be_updated` **do**

25: `new_score` ← UpdatedScore(*R*, `snps_covered_by_this_read`)

26: pq.ChangeScore(*R*, `new_score`)

27: **end for**

28: **end if**

29: **end if**

30: **end while**

31: **return** (`reads_in_slice, reads_violating_coverage`)

32: **end procedure**

### Algorithm 3 Bridging part of score based read selection

1: **procedure** Bridging(`reads_in_slice`, `undecided_reads`, `max_cov`)

2: *pq* ← ConstructPriorityqueue(`undecided_reads`)

3: `positions` ← ∪_*R*∈reads_in_slice_ `variants`(*R*) ∪ ∪_*R*∈undecided_reads_ `variants`(*R*)

4: `components` ← UnionFind(`positions`)

5: `bridge_reads` ← empty set

6: **for** read in `reads_in_slice` **do**

7: `components`. Merge(`variants(read)`)

8: **end for**

9: **while** *pq* not empty **do**

10: (`score, read`) ← pq.Pop

11: **if** |`components`.CoveredBy(variants(`read`))| ≥ 2 **then**

12: **if** adding read would not exceed `max_cov` for at least one position **then**

13: `bridge_reads`.Add(`read`)

14: `components`.Merge(variants(`read`))

15: **end if**

16: **end if**

17: **end while**

18: **return** `bridge_reads`

19: **end procedure**

WhatsHap and compared to the ground truth phasing. We examined the number of phased SNPs (phased), the number of unphased SNPs (unphased) the number of phased blocks (# blocks) and the number of correctly SNPs (true phased). Results are displayed in Table 1. Almost independent of the dataset the scoring-based read selection with the bridging surpasses the random approach in the number of correctly phased variants. Even without bridging, the scoring-based read selection provides an increased correctly phased variants compared to the random approach for all but one data set. The number of blocks in the scoring-based read selection with bridging is lower than the number of blocks in the random approach.

**Table 1:** Comparison of the random read selection with the score-based read selection approaches, one with (Scoring_bridging) and the other without bridging(Scoring_w/o_bridging) on a set of simulated datasets with a maximum coverage of both 5x and 15x.

| Dataset | Random |  |  |  | Scoring_bridging |  |  |  | Scoring_w/o_bridging |  |  |  |
| --- | --- | --- | --- | --- | --- | --- | --- | --- | --- | --- | --- | --- |
|  | Phased | Unphased | # blocks | True phased | Phased | Unphased | #blocks | True phased | Phased | Unphased | # blocks | True phased |
| A_1000 (15x) | 44 612 | 23 572 | 14 009 | 30 509 | 44 722 | 23 462 | 13 915 | 30 711 | 44 722 | 23 462 | 14 052 | 30 579 |
| A_10000 (15x) | 65 989 | 2 195 | 3 630 | 62 135 | 66 031 | 2 153 | 3 506 | 62 285 | 66 030 | 2 154 | 3 652 | 62 147 |
| A_5000 (15x) | 63 134 | 5 050 | 7 626 | 55 323 | 63 195 | 4 989 | 7 440 | 55 550 | 63 196 | 4 988 | 7 744 | 55 234 |
| A_50000 (15x) | 67 887 | 297 | 625 | 67 005 | 67 897 | 287 | 603 | 67 005 | 67 897 | 287 | 640 | 66 965 |
| Hiseq (15x) | 30 518 | 37 666 | 11 310 | 19 123 | 30 557 | 37 627 | 11 293 | 19 177 | 30 557 | 37 627 | 11 294 | 19 178 |
| Miseq 250 (15x) | 41 553 | 26 631 | 13 454 | 28 003 | 41 604 | 26 580 | 13 430 | 28 084 | 41 605 | 26 579 | 13 439 | 28 076 |
| Miseq 2500 (15x) | 39 681 | 28 503 | 12 706 | 26 871 | 40 002 | 28 182 | 12 465 | 27 428 | 40 001 | 28 183 | 12 985 | 26 910 |
| A_1000 (5x) | 44 612 | 23 572 | 14 009 | 30 509 | 44 721 | 23 463 | 13 915 | 30 702 | 44 721 | 23 463 | 14 273 | 30 348 |
| A_10000 (5x) | 65 989 | 2 195 | 3 630 | 62 135 | 66 027 | 2 157 | 3 506 | 62 286 | 66 028 | 2 156 | 4 094 | 61 702 |
| A_5000 (5x) | 63 134 | 5 050 | 7 626 | 55 323 | 63 195 | 4 989 | 7 440 | 55 564 | 63 196 | 4 988 | 8 291 | 54 720 |
| A_50000 (5x) | 67 887 | 297 | 625 | 67 005 | 67 709 | 475 | 682 | 66 721 | 67 710 | 474 | 968 | 66 441 |
| Hiseq (5x) | 30 144 | 38 040 | 11 516 | 18 519 | 30 557 | 37 627 | 11 293 | 19 174 | 30 556 | 37 628 | 11 364 | 19 107 |
| Miseq 250 (5x) | 41 553 | 26 631 | 13 454 | 28 003 | 41 602 | 26 582 | 13 430 | 28 068 | 41 601 | 26 583 | 13 662 | 27 838 |
| Miseq 2500 (5x) | 39 681 | 28 503 | 12 706 | 26 871 | 39 974 | 28 210 | 12 689 | 27 168 | 39 973 | 28 211 | 13 905 | 25 966 |

## 4 Discussion

As shown above, our novel score-based read selection provides some benefits in the connectivity and also in the increased number of phased or correctly phased variants. The overall quality has improved and the number of seleted reads under the same given coverage increased compared to the random approach. The algorithm described here has hence been integrated into the WhatsHap software.

We are currently comparing our heuristic approach to the flow-based algorithm proposed by Mäkinen *et al.* (2015). Our algorithm was designed to also work well when combining different types of reads (such as PacBio and Illumina mate pairs), which we plan to evaluate systematically in the future.

## References

Cilibrasi, R., Iersel, L. v., Kelk, S., and Tromp, J. (2007). The Complexity of the Single Individual SNP Haplotyping Problem. Algorithmica, 49(1), 13–36.

Cormen, T. H., Leiserson, C. E., Rivest, R. L., and Stein, C. (2009). Introduction to Algorithms. The MIT Press, third edition.

Glusman, G., Cox, H. C., and Roach, J. C. (2014). Whole-genome haplotyping approaches and genomic medicine. Genome Medicine, 6(9), 73.

He, D., Choi, A., Pipatsrisawat, K., Darwiche, A., and Eskin, E. (2010). Optimal algorithms for haplotype assembly from whole-genome sequence data. Bioinformatics, 26(12), i183–i190.

Mäkinen, V., Staneva, V., Tomescu, A. I., and Valenzuela, D. (2015). Interval scheduling maximizing minimum coverage. arxiv, 1508.07820.

Patterson, M., Marschall, T., Pisanti, N., van Iersel, L., Stougie, L., Klau, G. W., and Schönhuth, A. (2015). WhatsHap: Weighted haplotype assembly for future-generation sequencing reads. Journal of Computational Biology, 22(6), 498–509.

Tewhey, R., Bansal, V., Torkamani, A., Topol, E. J., and Schork, N. J. (2011). The importance of phase information for human genomics. Nature Reviews Genetics, 12(3), 215–223.

